# BrainNET: Inference of brain network topology using Machine Learning

**DOI:** 10.1101/776641

**Authors:** Gowtham Krishnan Murugesan, Chandan Ganesh, Sahil Nalawade, Elizabeth M Davenport, Ben Wagner, Kim Won Hwa, Joseph A. Maldjian

## Abstract

**Objective:** To develop a new fMRI network inference method, BrainNET, that utilizes an efficient machine learning algorithm to quantify contributions of various regions of interests (ROIs) in the brain to a specific ROI.

**Methods:** BrainNET is based on Extremely Randomized Trees (ERT) to estimate network topology from fMRI data and modified to generate an adjacency matrix representing brain network topology, without reliance on arbitrary thresholds. Open source simulated fMRI data of fifty subjects in twenty-eight different simulations under various confounding conditions with known ground truth was used to validate the method. Performance was compared with correlation and partial correlation (PC). The real-world performance was then evaluated in a publicly available Attention-deficit/hyperactivity disorder (ADHD) dataset including 134 Typically Developing Children (mean age: 12.03, males: 83), 75 ADHD Inattentive (mean age: 11.46, males: 56) and 93 ADHD Combined (mean age: 11.86, males: 77) subjects. Network topologies in ADHD were inferred using BrainNET, correlation, and PC. Graph metrics were extracted to determine differences between the ADHD groups.

**Results:** BrainNET demonstrated excellent performance across all simulations and varying confounders in identifying true presence of connections. In the ADHD dataset, BrainNET was able to identify significant changes (p< 0.05) in graph metrics between groups. No significant changes in graph metrics between ADHD groups was identified using correlation and PC.

## 1. Introduction

The brain is a complex interconnected network that balances segregation and specialization of function with strong integration between regions, resulting in complex and precisely coordinated dynamics across multiple spatiotemporal scales [1]. Connectomics and graph theory offer powerful tools for mapping, tracking, and predicting patterns of disease in brain disorders through modeling brain function as complex networks [2]. Studying brain network organization provides insight in understanding global network connectivity abnormalities in neurological and psychiatric disorders [3]. Several studies suggest that pathology accumulates in highly connected hub areas of the brain [4, 5] and that cognitive sequelae are closely related to the connection topology of the affected regions [6]. An understanding of network topology may allow prediction of expected levels of impairment, determination of recovery following an insult and selection of individually tailored interventions for maximizing therapeutic success [7]. A large number of network inference methods are being used to model brain network topology with varying degrees of validation. A recent study [8] evaluated some of the most common methods, including correlation, partial correlation, and Bayes NET, to infer network topology using simulated resting state functional magnetic resonance images (fMRI) data with known ground truth and found that performance can vary widely under different conditions.

Development of statistical techniques for valid inferences on disease-specific group differences in brain network topology is an active area of research. Machine learning methods have been used in neuroimaging for disease diagnosis and anatomic segmentation [9, 10]. Deep learning and classical machine learning methods are used to distinguish diseased groups from controls based on features extracted from fMRI [11, 12]. Very few studies have attempted to apply machine learning methods on direct time series of fmri to infer brain networks [9, 13-15]. Recent work in machine learning approaches for inference of Gene Regulatory Networks (GRN) has demonstrated excellent performance [16-18]. Interestingly, these same approaches to gene regulatory networks can be used to infer brain networks. In this study, we describe a new network inference method called BrainNET, inspired by machine learning methods used to infer GRN [19].

Validation of BrainNET was performed using fMRI simulations with known ground, as well as in real-world ADHD fMRI datasets. In this study, publicly available resting state fMRI simulated data [8] was used to validate BrainNET’s ability to infer networks. The real-world performance of BrainNET was then evaluated in a publicly available data set of Attention-deficit/hyperactivity disorder (ADHD). ADHD is one of the most common neurodevelopmental disorders in children with significant socioeconomic and psychological effects [20, 21] and it is very difficult to diagnose [22]. ADHD has widespread but often subtle alterations in multiple brain regions affecting brain function [23, 24]. Neuro Bureau, a collaborative neuroscience forum, has released fully processed open source fMRI data “ADHD-200 preprocessed” from several sites [25, 26] providing an ideal dataset to test the BrainNET model and compare its performance with standard correlation and partial correlation (PC), which is the most widely used methodology to infer brain networks using fMRI data.

## 2. Materials and Methods

### 2.1. Datasets

#### 2.1.1. MRI Simulation Data

Open source rs-fMRI simulation data representing brain dynamics was used to validate the BrainNET model [8]. The data were simulated based upon the dynamic causal modeling fMRI forward model, which uses the non-linear balloon model for vascular dynamics, in combination with a neural network model [8]. The open source dataset has 28 simulations; each including simulated data for 50 subjects with a varying number of nodes and several confounders (e.g., shared input between the nodes, varying fMRI session lengths, noise, cyclic connections and hemodynamic lag variability changes). Additional details on the simulations can be found in the original study [8] (supplementary table I).

#### 2.1.2. ADHD data

Preprocessed rs-fMRI data were obtained from the ADHD-200 database (http://fcon1000.projects.nitrc.org/indi/adhd200/). Seven different sites contributed to the ADHD-200 database for 776 rs-fMRI data acquisitions. The data were preprocessed using the Athena pipeline and was provided in 3D NifTI format. Additional information on the Athena pipeline and “ADHD 200 preprocessed” data is detailed by Bellec et al [25].

In our study, subjects identified with ‘No Naïve medication’ status, or questionable quality on rs-fMRI data were excluded. The remaining subjects were age-matched between the groups resulting in 135 Typically Developing Children (TDC) (mean age: 12.00, males: 83), 75 ADHD Inattentive (ADHD-I) (mean age: 11.46, males: 56) and 93 ADHD Combined (ADHD-C) (mean age: 11.86, males: 77) subjects. Mean time series from 116 ROI’s in the AAL atlas [27] were extracted using the NILEARN package [28].

### 2.2. BrainNET Model Development

The objective of BrainNET is to infer the connectivity from fMRI data as a network with N different nodes in the brain (i.e., ROI’s), where edges between the nodes represent the true functional connectivity between nodes. At each node, there are measurements from m time points *X* = {*x*_1_,*x*_2_,*x*_3_, *x*_4_, ….,*x*_*N*_}, where x_i_ is the vector representation of m time points measured as

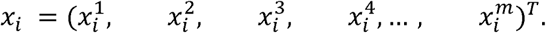

Our method assumes that fMRI measurement of BOLD (Blood Oxygen Level Dependent) activation at each node is a function of each of the other nodes’ activation with additional random noise.

For the j^th^ node with m time points, a vector can be defined denoting all nodes except the j^th^ node as *x*_−*j*_ = (*x*_1_,*x*_2_,*x*_*j*−1_,*x*_*j*+1_,….,*x*_*N*_), then the measurements at the j^th^ node can be represented as a function of other nodes as

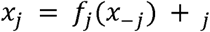

where □_j_ is random noise specific to each node_j_. We further assume that function *f*_j_ () only exploits the data of nodes in x_-j_ that are connected to node_j_. The function *f*_j_ () can be solved in various ways in the context of machine learning. Since the nature of the relationship between different ROIs in the brain is unknown and expected to be non-linear [29], we choose a tree based ensemble method as it works well with a large number of features with non-linear relationships and is computationally efficient. We utilized Extremely Randomized Trees (ERT), an ensemble algorithm similar to Random Forest, which aggregates several weak learners to form a robust model. ERT uses a random subset of predictors to select divergences in a tree node and then selects the “best split” from this limited number of choices [30]. Finally, outputs from individual trees are averaged to obtain the best overall model [31]. BrainNET infers a network with N different nodes by dividing the problem into N different sub problems, and solving the function *f*_j_ () for each node independently as illustrated in Figure 1. The steps are listed below: For j = 1 to N nodes

**Figure 1:**
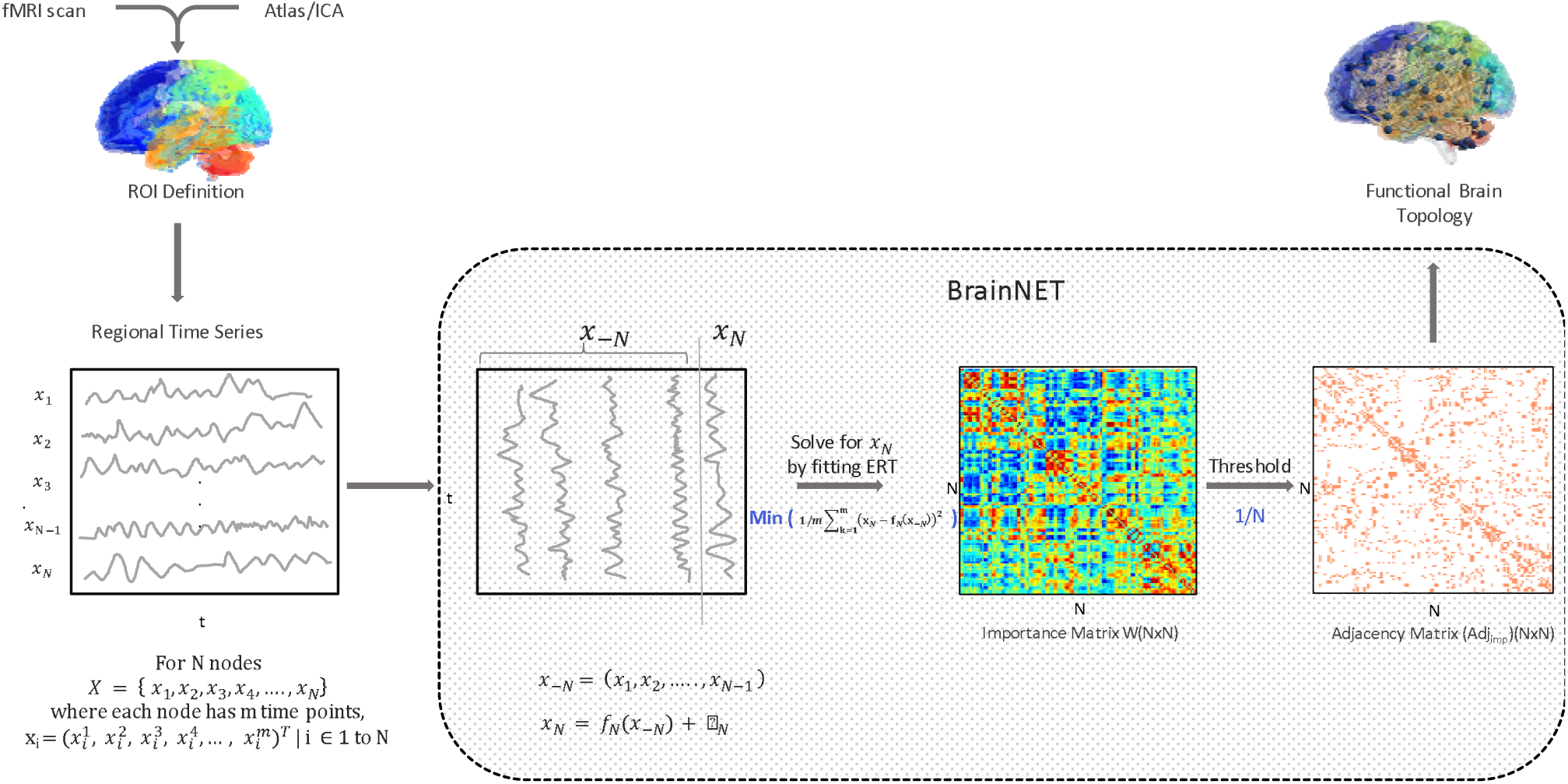
Schematic overview of the BrainNET model. For N nodes in fMRI data (X), each node will have m time points such that X = {x_1_,x_2_,x_3_, x_4_, ….,x_N_}, where x_i_ is the vector representation of m time points measured as 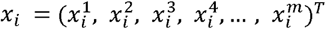. Each node’s time series (x_n_) is predicted from all other nodes time series (x_-n_) using the ERT regressor. Node Importance of each node for predicting the target node are extracted and populated in the importance matrix. The average of the upper and lower triangle of the matrix is thresholded at (1/Num of Nodes) to obtain an adjacency matrix representing the network topology.

- Fit the ERT regressor with all the nodes data, except the j^th^ node, to find the function f_j_ that minimizes the following mean squared error:

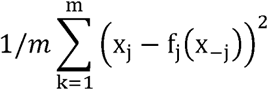
- Extract the weight of each node to predict node j,

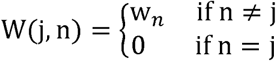

where w_n_ is the weight of node to predict node j and n= 1 to N.
- Append the weights values to the Importance matrix

The importance score for each node (Node_j_) to predict (Node_i_) is defined as the total decrease in impurity due to splitting the samples based on Node_j_ [30]. Let “S” denote a node split in the tree ensemble and let (S_L_, S_R_) denote it’s left and right children nodes. Then, the decrease in impurity ΔImpurity(S) from node split “S” based on Node_j_ to predict Node_i_ is defined as

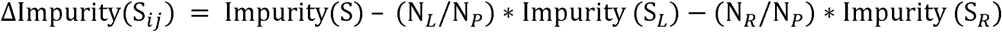

where, S_L_ and S_R_ are left and right splits and N_P_, N_L_, N_R_ are number of samples reaching parent, left and right nodes respectively. Let 𝕍k be the number of ensembles, which uses ROI_j_ for splitting trees. Then, the importance score for Node_j_ for predicting Node_i_ is calculated as the average of node impurities across all trees, i.e. Importance of ROI_ji_

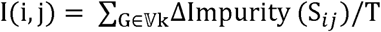

where T is the number of trees in the ensemble.

Importance values extracted using a typical Random Forest model can be biased in the presence of two or more correlated features since the model will randomly assign importance to any one of the equally important features without any preference [32]. This problem is avoided by using the ERT regressor.

The importance of each node to predict all other node time series is extracted from the model and an NxN (where N is the number of nodes) importance matrix is generated with the diagonal equal to zero. Each row of the importance matrix represents normalized weights of each node in predicting the target node. The extracted adjacency matrix is affected in two ways. First, due to the row-wise normalization, the upper triangular values of the importance matrix are not same as the lower triangle values. We therefore take the average of the upper triangle and the lower triangle of the matrix to make it symmetric to determine the presence of connection between the nodes. This procedure does not allow directionality of the connections to be determined. Second, again because of the row-wise normalization, the sum of each row in the importance matrix is one. Since the importance values are normalized with respect to number of nodes in the analysis, we used a threshold equal to a theoretical probability value that is inversely proportional to the number of nodes (i.e., threshold = 1/number of nodes) in the network to produce a final adjacency matrix representing the network topology. This results in a dynamically changing threshold based on the number of nodes in the network.

### 2.3. Analysis

#### 2.3.1. Evaluation of inference methods on simulation data

##### 2.3.1.1. Evaluation of inference methods on simulation data using C-sensitivity

The network topology was inferred using BrainNET, correlation and PC. The network topology inferred by correlation and PC method may vary drastically based on the values used to threshold connectivity matrix. Hence, we evaluated the ability of the inference methods based on BrainNET, correlation and PC to detect the presence of connection between the nodes in terms of c-sensitivity. C-sensitivity quantifies how well the true positives (TP) are separated from the false positives (FP) by measuring the fraction of TPs that are estimated with a higher connection strength than the 95th percentile of the FP distribution. C-sensitivity is a measure of success in separating true connections from false positive connections and it is calculated by counting number of true positive above 95th percentile of false positives and then divided by total number of true positives [8].

Effects of simulation parameters such as TR (repetition time), number of nodes, noise, HRF (Hemodynamic Response Function) standard deviation, shared inputs, bad ROI’s (Region of Interest), backward, strong and cyclic connections and strong inputs on c-sensitivity of the inference methods were evaluated using multiple linear regression. For each inference method, and for each subject, a multiple regression was fit to the 28 simulations c-sensitivity values, with the different simulation parameters as model covariates. The model fitting was carried out separately for each of the 50 subjects and the parameter estimates from each regression were then summarized across subjects in terms of their effect size (mean/standard deviation) [8].

##### 2.3.1.2. Evaluation of inference methods on simulation data using threshold

Thresholding can be applied to suppress spurious connections that may arise from measurement noise and imperfect connectome reconstruction techniques and to potentially improve statistical power and interpretability [7]. However, based on the threshold value, the connection density of each network inferred by correlation and PC may vary from network to network after the threshold has been applied. Using a less stringent lower threshold values results higher false positive values (lower sensitivity) and more stringent threshold results in higher false negatives(lower specificity). This can lead to wide variability in computed graph metrics, as they are typically susceptible to the number of edges in a graph. Identifying an appropriate threshold to infer the underlying brain network topology is critical.

Hence, we evaluated the specificity, sensitivity and accuracy of correlation and PC under varying thresholds. The results shows that the network topology inferred using correlation and PC method may vary drastically based on the threshold values (Fig.2). An optimum threshold for correlation methods can be very difficult to find in real life experimental data. However, given the ground truth for the simulation data we calculated the optimum threshold values for the correlation methods and compared their performance at optimum threshold with BrainNET. It is important to note that BrainNET is not optimized on this simulation data and the threshold is based on the number of nodes inferred in the network. Specificity, sensitivity and accuracy of correlation and PC at threshold values of 30% (Corr_30_, PC_30_) and optimum (Corr_opt_, PC_opt_) values are estimated and compared with BrainNET. We further evaluated the specificity, sensitivity and accuracy of the Corr_opt_ and PC_opt_ with BrainNET for each simulation.

**Figure 2:**
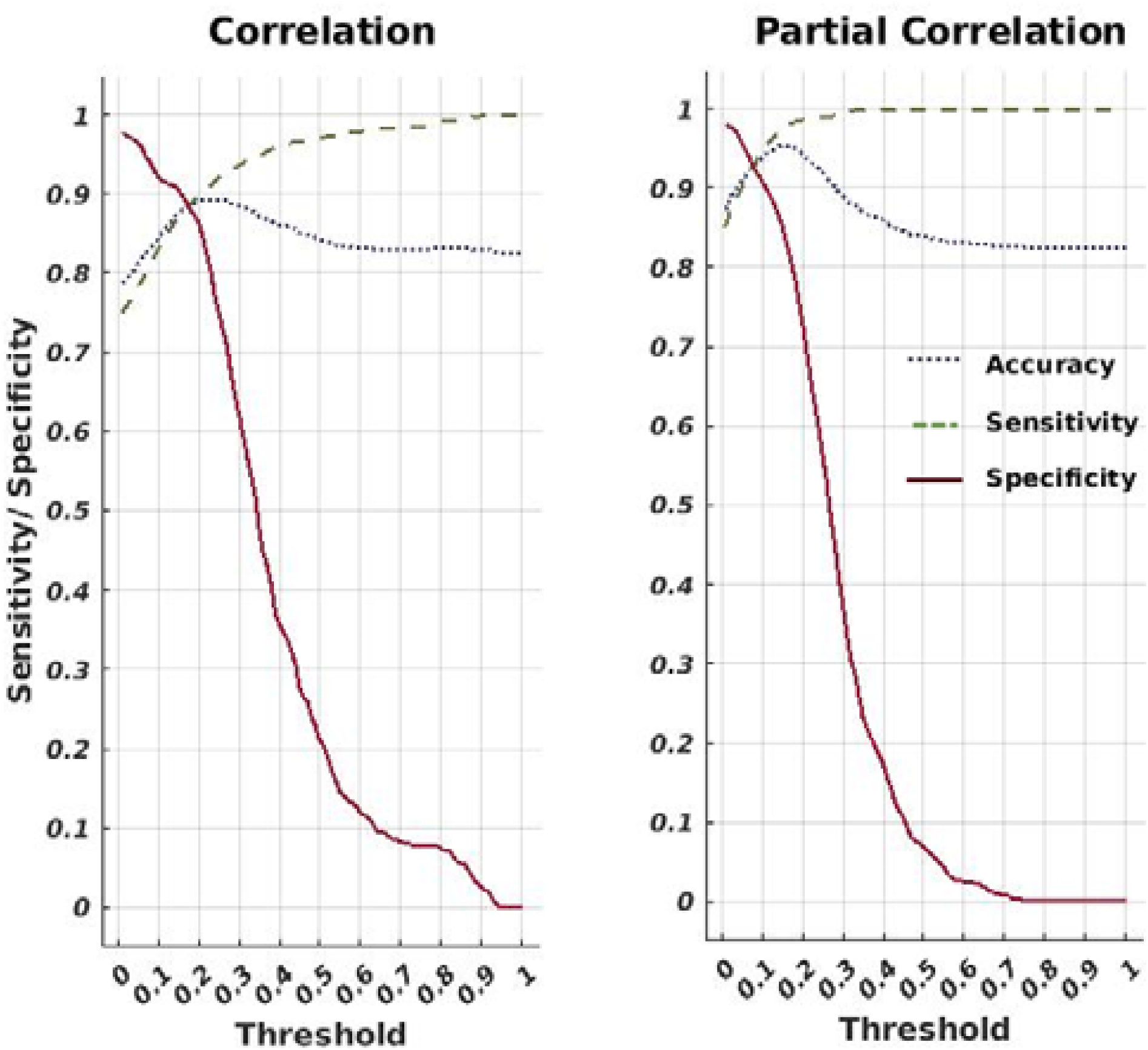
Sensitivity analysis for correlation and partial correlation. Average sensitivity (true positive rate) and specificity across 28 simulations for the correlation and PC method is plotted as a function of threshold ranging between zero to one hundred percent. Optimum threshold is found using simulation ground truth at 20% and 16% for correlation and PC respectively.

##### 2.3.1.3. Evaluation of inference methods on ADHD data

BrainNET, correlation and PC was applied to the real-world ADHD data to evaluate whole brain network changes in ADHD subtypes (i.e., ADHD-Combined (ADHD-C), ADHD Inattentive (ADHD-I) compared to Typically Developing Children (TDC). Mean time series from 116 ROI’s in the AAL atlas [27] were extracted using the NILEARN package [28]. The BrainNET model was applied to extract an importance matrix for each subject. The importance matrix was then thresholded at 1/number of nodes (e.g., 1/116 for the AAL atlas regions) to obtain an adjacency matrix for each subject (BN). Functional Network connectivity was calculated between the 116 ROIs using correlation and PC. The connectivity matrices are thresholded at a threshold of 20% and 30% (Corr_20_, Corr_30_, PC_20_, and PC30) (No optimum threshold for real world experimental data). Graph theoretic metrics were extracted using each of these methods for each group. Network differences between the three groups TDC, ADHD-I and ADHD-A were then computed using t-tests on the graph metrics. Site effects and effects of age and handedness were removed using the Combat multi-site harmonization method[33], an effective harmonization technique that removes both unwanted variation associated with the site and preserves biological associations in the data [34].

###### Graph Metrics

Graph theoretical metrics representing global and local characteristics of network topology were used to compare between the groups in the ADHD dataset. The GRETNA MATLAB toolbox (v2.0,https://www.nitrc.org/projects/gretna/) was used to extract additional graph theoretical metrics including shortest path length, global network efficiency, and betweenness centrality[35]. The Networkx package in python was used to extract the graph theoretical metrics, including Density, Average Clustering Coefficient and Characteristic Path length [36]. Two sample t-tests between groups were performed using the GRETNA toolbox. Bonferroni multiple comparisons correction was applied with statistical significance set at p < .05.

###### Node Metrics

The **Nodal Shortest Path Length** (NSPL) is defined as the shortest mean distance from a particular node to all other nodes in the graph. Shorter NSPL represents greater integration[1]. The **Betweenness Centrality (BC)** measures a node’s influences in information flow between all other nodes [37]. BC quantifies the influence of a node and is defined as the number of shortest paths passing through it.

###### Global Metrics

**Network Efficiency** is a more biologically relevant measure representing the ability of the network to transmit information globally and locally. Networks with high efficiency, both globally and locally, are said to be economic small world networks. [38]. The **density** of the graph is defined as the ratio of number of connections in the network to the number of possible connections in the network. **Average Clustering** is the fraction of a node’s neighbors that are neighbors of each other. The clustering coefficient of a graph is the **average clustering coefficien**t (ACC) over all nodes in the network. Networks with high clustering coefficient are considered locally efficient networks. **Characteristic Path length** (CPL) is the average shortest path length between nodes in the graph, with a minimum number of edges that must be traversed to get from one node to another. CPL indicates how easily information can be transferred across the network [1].

## 3. Experimental Results

### 3.1. Simulation Data

#### 3.1.1. Evaluation of inference methods using C-sensitivity

BrainNET performed significantly better than correlation (p<0.001) and equivalent to partial correlation (p > 0.05) methods with c-sensitivity of 79.53%, 59.82% and 75.75% respectively across 28 simulations (Fig.3A) (supplementary table.II). The study on effect of simulation parameters on the c-sensitivity of inference methods shows that increasing the number of nodes and session duration has strong positive effect on correlation c-sensitivity but not much on BrainNET and PC (Fig.3B). Selection of TR has stronger effect on correlation than BrainNET and PC method. Presence of noise in the signals affected the BrainNET the least compared to other methods. Having shared inputs between the nodes affected the correlation and PC method more drastically than BrainNET method. Selection of bad ROI’s with mixed time series between them affects all the inference methods negatively, however selection of bad ROI’s with randomly random time series mixed between them doesn’t affect the inference methods drastically. Presence of cyclic and backward connection between the nodes affected the correlation and PC methods but not BrainNET. Having more connections and stationary-nonstationary connections between the nodes affected the PC and correlation method more than the BrainNET method. The presence of only one strong input affected the performance of BrainNET method but not the other methods. In summary, BrainNET performance is robust under various confounding factors but prone to selection of inaccurate ROI’s with mixed time series between them and networks with only one strong input. Both PC and correlation methods are affected by shared inputs between the nodes, selection of inaccurate ROI’s, backward, cyclic and stationary-nonstationary connections between the nodes.

**Figure 3:**
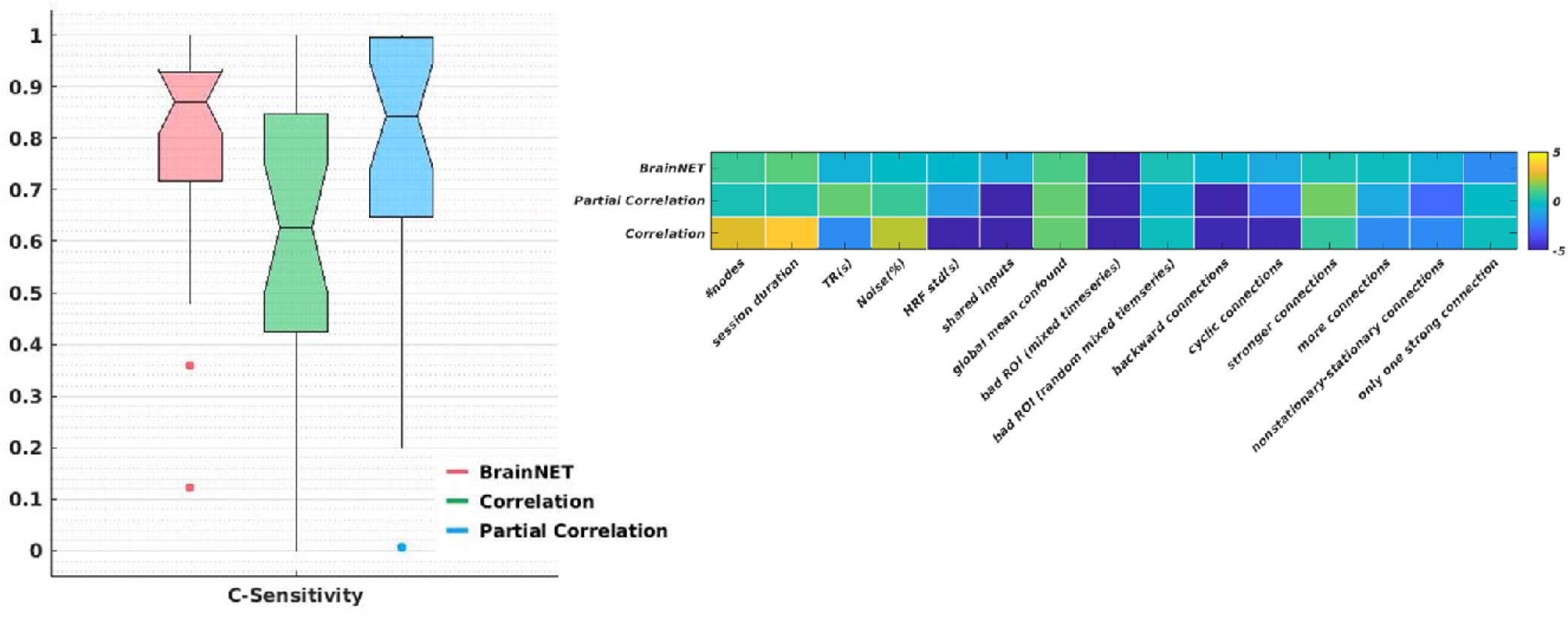
C-sensitivity. Boxplots of c-sensitivity for BrainNET, correlation and partial correlation (PC) (left). The effects of different simulation parameters on the c-sensitivity of inference methods using multiple linear regression. The colorbar represents effect size of each simulation parameter on the c-sensitivity of inference methods.

#### 3.1.2. Evaluation of inference methods using threshold

The accuracy, sensitivity, and specificity for each method across all 28 simulated data sets are estimated at thresholds of 30% and at optimum value for correlation and PC, and at threshold of 1/number of nodes for BrainNET (Fig.4). BrainNET achieved higher accuracy and specificity at threshold at 30% compared to the Corr_30_ and PC_30_ method as shown in Fig.4. PCopt achieved slightly higher accuracy than BrainNET across 28 simulations, but no significant difference in terms of specificity and accuracy even at it optimum (p>0.05) (Fig.5). As expected the specificity and sensitivity of correlation and PC methods varies with threshold and it will be difficult to find an optimum threshold in a real life dataset. BrainNET showed more robust performance with little variance across the simulation compared to other methods.

**Figure 4:**
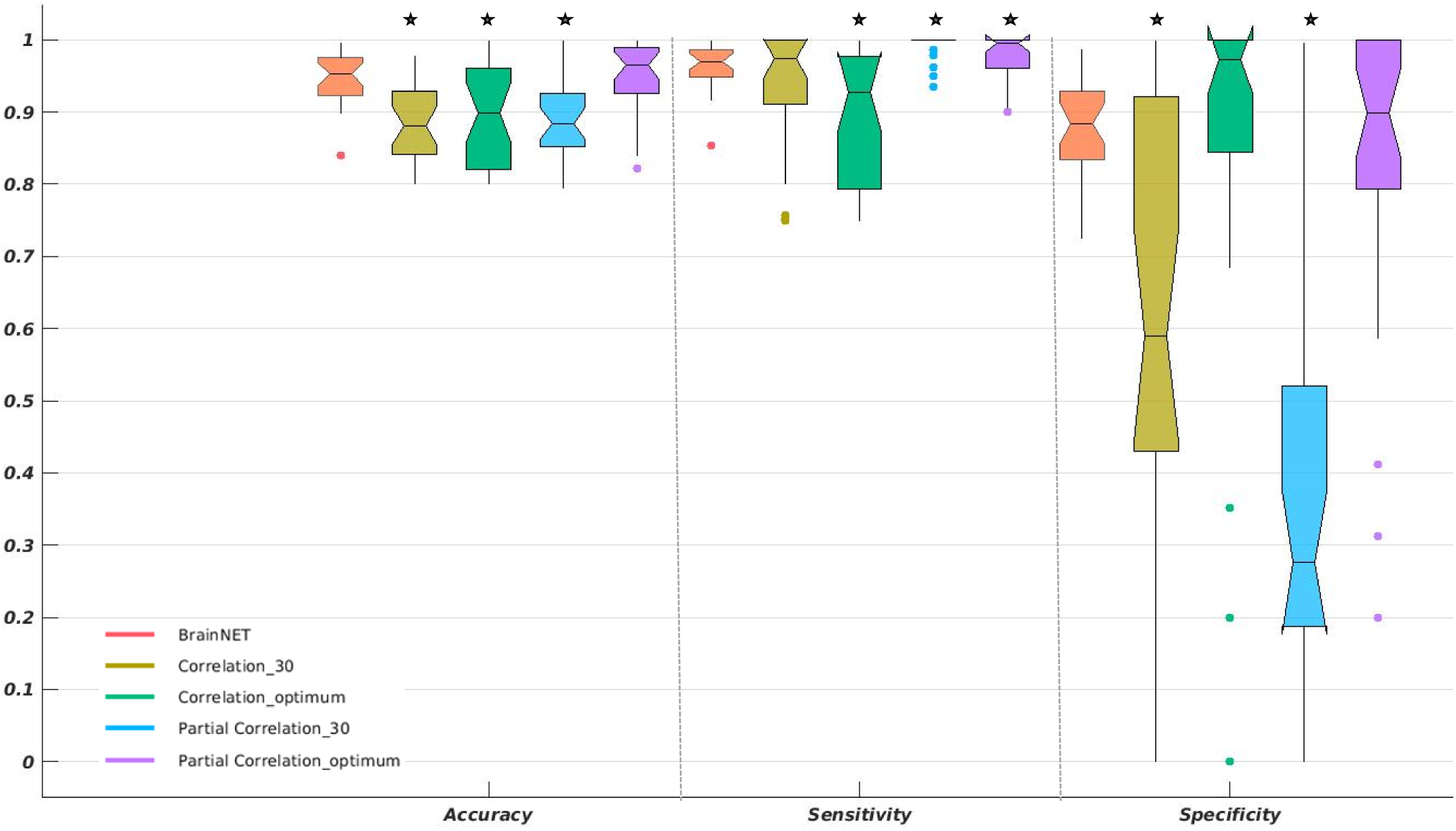
Evaluation of inference methods under varying thresholds. Boxplots of Accuracy (Left), Sensitivity (Middle) and Specificity (Right) across 28 simulations for correlation and PC for optimum and thirty percent threshold (Corr_opt_, Corr30, PC_opt_ and PC_30_), and BrainNET. ‘*’ represents statistically significant differences from BrainNET performance.

**Figure 5:**
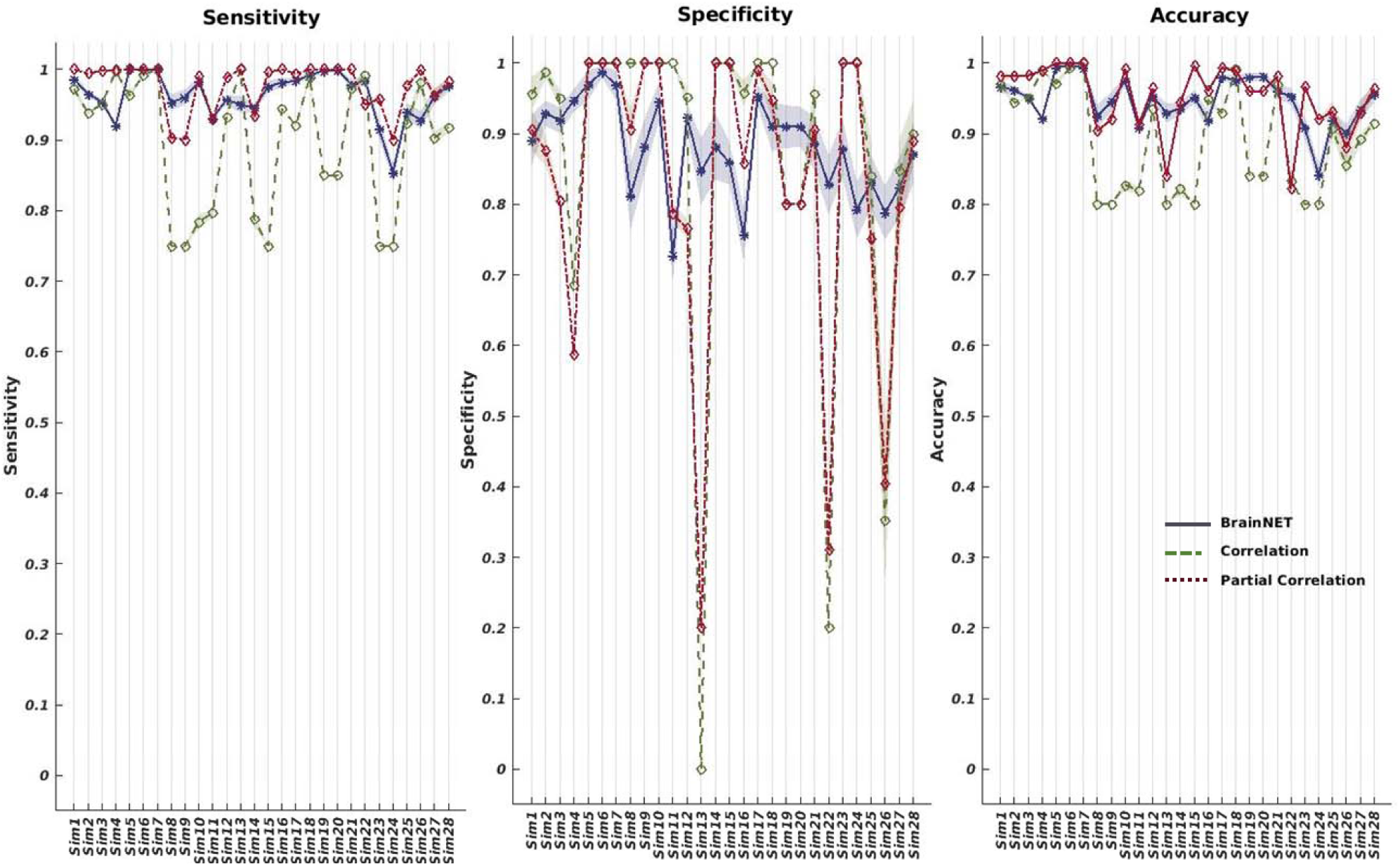
Comparison of Correlation (Corr_opt_) and partial correlation (PC_opt_) at their optimum threshold to BrainNET. Accuracy (Left), Sensitivity (Middle) and Specificity (Right) for correlation, BrainNET and PC for 28 simulations. Sensitivity, specificity and accuracy are all robust across different simulation cases, while PC and correlation methods show fluctuations even with their optimal threshold for functional connectivity.

##### 3.1.2.1. Evaluation of inference methods on ADHD data

BrainNET was able to identify significant changes (p < 0.05) in global network efficiency, network density, characteristic path length, betweenness centrality and shortest path in the ADHD data. Correlation and PC was not able to detect significant changes in any of the whole-brain analyses (Corr_20_, Corr_30_, PC_20_, and PC_30_).

###### TDC and ADHD

Statistical analysis of the BrainNET adjacency matrix demonstrated a significant decrease in global Network efficiency, an increase in CPL and an increase in the shortest path length in the right medial temporal gyrus in ADHD compared to TDC (Fig.7A). While the analysis of the correlation adjacency matrix did not show any significant changes, the PC_30_ demonstrated a trending increase in CPL in ADHD compared to TDC (p=0.07). Betweenness centrality and Node level local efficiency did not show any changes between the groups in any of the three methods.

###### TDC and ADHD-I

Statistical analysis of the BrainNET adjacency matrix demonstrated a significant decrease in global network efficiency, a decrease in density, an increase in CPL and an increase in shortest path length in the right superior orbital right, right heschl’s gyrus and right medial temporal gyrus nodes in the ADHD-I group compared to TDC (Fig.7A). The correlation method did not show any relation between the groups. No relationship was found in other graph metrics studied in any of the methods.

###### TDC and ADHD-C

Statistical analysis of the BrainNET adjacency matrix demonstrated a significant decrease in density. No significant relation was found in any other graph metrics for any of the three methods.

###### ADHD-I and ADHD-C

Statistical analysis of the BrainNET adjacency matrix demonstrated a significant decrease in global network efficiency, a decrease in density, an increase in CPL, an increase in shortest path length of the right olfactory node (Fig.7A). A significant increase in betweenness centrality of the right precuneus node in the ADHD-I group compared to ADHD-C was observed for both BrainNET (Fig.7B). No relationship was found in other graph metrics studied for any of the methods.

## 4. Discussion

BrainNET was developed to infer brain network topology using ERT [39]. The ERT regressor is used to develop a tree based ensemble model to predict each node’s time series from all other node time series. The tree based ensemble methods are ideal for inferring complex functional brain networks as they are efficient in learning non-linear patterns even where there are a large number of features [40]. The importance matrix is then thresholded to generate an adjacency matrix representing the fMRI topology. The BrainNET model is applicable to both resting-state and task-based fMRI network analysis. It can be easily adapted to datasets with varying session lengths and can be used with different parcellation schemes. A unique feature of the BrainNET approach is that it is implemented at the subject level. It does not need to be trained on big datasets as it infers the network topology based on each individual subject’s data.

### 4.1. BrainNET Inference of Network Topology in Simulated fMRI Data

#### 4.1.1. Evaluation of inference methods using C-sensitivity

BrainNET demonstrated excellent performance across all the simulations and varying confounders. It achieved significantly higher c-sensitivity than correlation (p<0.05) and equivalent to PC (p=0.38) (Fig.2A). BrainNET performance remained high in the simulations across varying session lengths, number of nodes, neural lags, cyclic connections, and changing number of connections. BrainNET performed weakest in simulations with one primary strong external source around the network. This causes every node to be highly correlated with other nodes and it becomes very difficult to distinguish direct from indirect connections [8]. It is important to highlight that this kind of one strong external input just for one node is highly unlikely in real life scenarios. BrainNET, similar to PC and correlation methods, was affected by selection of bad ROI’s with time series mixed between them. In this simulation, there are 10 nodes, and each node shares a relatively small amount of the other node time series in a proportion of 0.8:0.2. Since the features have shared data between the nodes in this simulation, it limits discrimination of true connectivity between nodes. The leakage of data between nodes can be minimized in fMRI analysis by selecting independent regions using functionally derived parcellation or methods such as ICA.

Our choice of threshold for BrainNET [1/(number of nodes)], represents a theoretical probability for the presence of a connection between the nodes. One concern with this approach is that as the number of nodes increases, the threshold similarly decreases, and may result in increased false positives at this low threshold value. The study on effect of number of nodes on the c-sensitivity of BrainNET shows that c-sensitivity of BrainNET increases with increasing the number of nodes (Fig.2B). This can be interpreted that the ability of BrainNET to distinguish between true and false positives increases with the increasing number of nodes and the corresponding lower threshold values doesn’t necessarily affect its inference.

Shared inputs between the nodes, can be thought of as distinct sensory inputs that feed into one or more nodes. These shared inputs between the nodes could be deleterious if not modelled [8]. BrainNET is robust to the shared inputs between the nodes, whereas c-sensitivity of PC and correlation are negatively affected (Fig.2B). The performance on varying connection strength over time was tested by simulations of stationary-nonstationary connection strengths between the nodes. BrainNET was least affected by nonstationary connection strengths between the nodes (Fig.2B). The robust performance of BrainNET in simulations with increasing number of nodes, TR, shared inputs, backward, cyclic and non-stationary connections represents a promising aspect of the BrainNET method for inferring brain network topology in real life experimental data (Fig.2B).

#### 4.1.2. Evaluation of inference methods using thresholds

In this study we compared the performance of correlation and PC in inferring underlying network topology at optimum threshold values estimated using ground truth (Corr_opt_ and PC_opt_). We also compared the performance of these methods against BrainNET at 30% threshold (Corr_30_ and PC_30_) (Fig.4). BrainNET performed significantly better than PC_30_, Corr_30_ and Corr_opt_ (p<0.05). At its optimum threshold, PC_opt_ performed relatively equivalent to BrainNET in terms of accuracy. However, PC_opt_ showed decreased specificity with increasing number of nodes (sim1-4) and presence of nonstationary, backward connections (sim13 and sim22) (Fig.5). Nonstationary connections represent the varying strengths of connections between nodes which are believed similar to those at the neuronal level and being studied in fMRI. The higher sensitivity and lower specificity of PC_opt_ represents higher numbers of false positive connections which will affect the statistical power of group analysis. The results show that the performance of the PC and correlation method vary under different thresholds and that BrainNET had better performance than these methods even in their optimum (PC_opt_ and Corr_opt_) (Fig.4).

A major strength of the BrainNET approach is that it provides a unique threshold to determine the true network topology. In correlation-based approaches, there is no defined correlation cutoff to determine the true network topology. Instead, multiple approaches are employed, or multiple thresholds applied to generate different networks. Typically, the network cost has been used to define the cutoff value for defining true connections in correlation-based approaches [41]. Multiple costs are then applied to generate multiple instances of the network topology, and analyses are performed to determine the variation in network metrics across these costs, or variation in group differences across thresholds [42]. The BrainNET approach provides a single threshold obviating the need for these imprecise and convoluted thresholding approaches.

### 4.2. Evaluation of inference methods on ADHD Data

#### Global metrics

Previous studies have shown that ADHD is often associated with changes in functional organization of the brain and lower network efficiencies in ADHD [21, 24]. BrainNET was effective in identifying the subtle changes in the ADHD subjects and supports the notion that the functional organization of brain changes in ADHD (Fig.6) by identifying statistically significant changes in graph metrics between ADHD subjects and typically developing children (Table 2).

**Figure 6:**
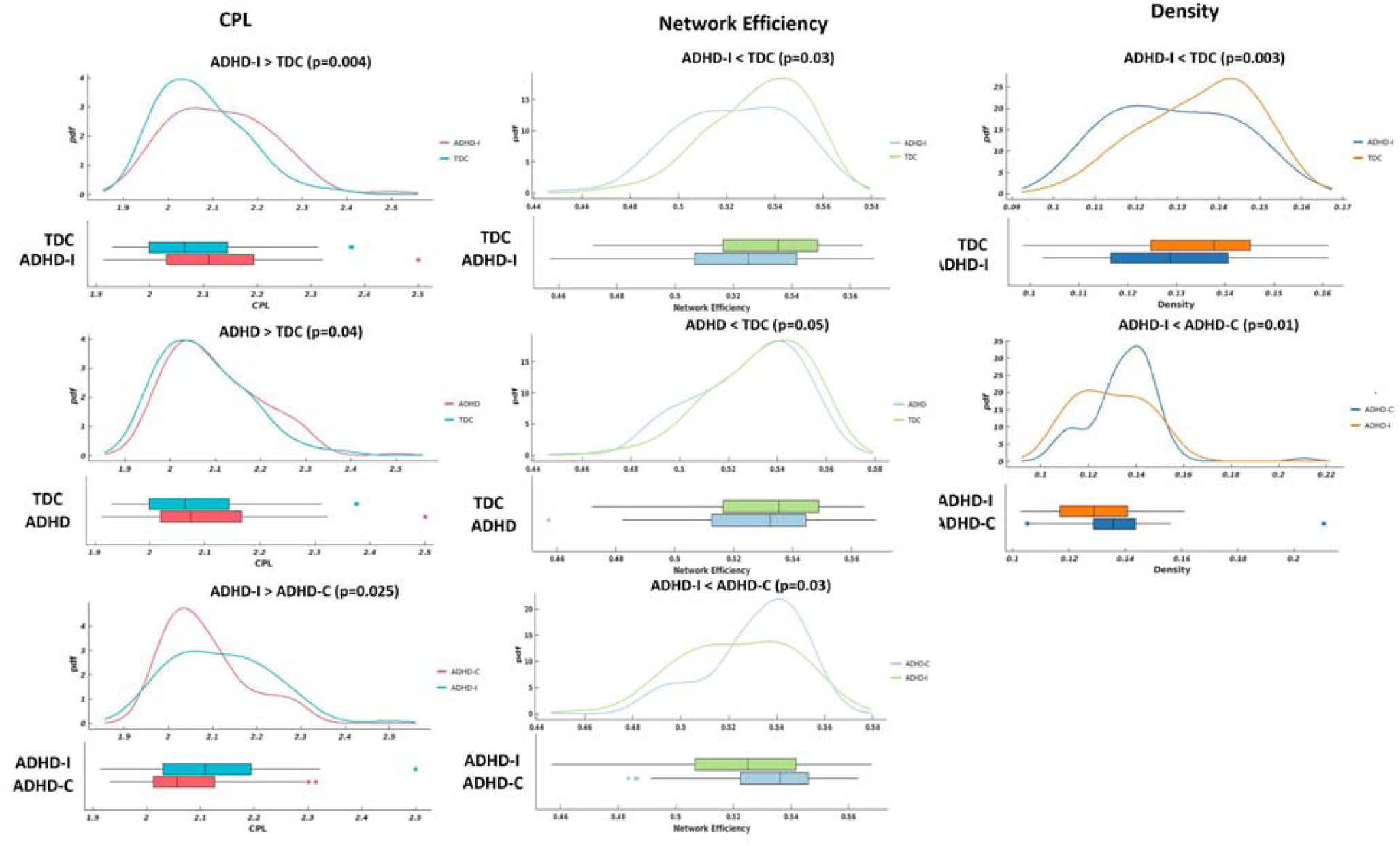
Global graph metrics. The probability density functions and boxplots of global graph metrics with significant changes (p<0.05) between the groups, ADHD (both ADHD-I and ADHD-C), ADHD-I, ADHD-C and TDC. CPL – Characteristic Path Length.

Our results demonstrate that there is a decrease in density, network efficiency and an increase in CPL in ADHD compared to TDC. A decrease in density suggests that the number of connections is decreased in ADHD compared to TDC. This can be interpreted as an increase in the cost of wiring in the brain. The increase in CPL and decrease in network efficiency is expected given that there is a decrease in density suggesting that there is increased difficulty in transferring information across the brain in ADHD. The observed abnormalities in global network topology was identified in ADHD-I but not in participants with ADHD-C compared with TDC, however changes between the ADHD-C and ADHD-I were observed. The differential changes observed between the ADHD subtypes may reflect clinical distinctions between the inattentive and combined subtypes of ADHD. Further investigations may shed light on detailed brain-behavior phenotype associations in this neuropsychiatric disorder [43, 44].

#### Local Metrics

BrainNET identified increased NSPL in ADHD-I compared to ADHD-C suggesting lesser integration of the prefrontal cortex (PFC) in ADHD-I. The PFC is a part of Default Mode Network (DMN) and plays a crucial role in regulating attention, behavior, and emotion, with the right hemisphere specialized for behavioral inhibition [45]. The DMN refers to the brain circuitry that includes the medial prefrontal cortex, posterior cingulate, precuneus, and the medial, lateral, and inferior parietal cortices [47]. These results support previous studies demonstrating that ADHD is associated with structural changes and decreased function of the PFC circuits, especially in the right hemisphere [45]. BrainNET also demonstrated that betweenness centrality of the right precuneus, also a part of DMN was increased in ADHD-I compared to ADHD-C group. This suggests increased influence of the precuneus in ADHD-I (Fig.7B). Abnormalities within the DMN have also been found in children in previous studies with ADHD and especially changes in centrality of the right precuneus, which is an important discriminatory feature for classifying ADHD-I and ADHD-C [46].

**Figure 7:**
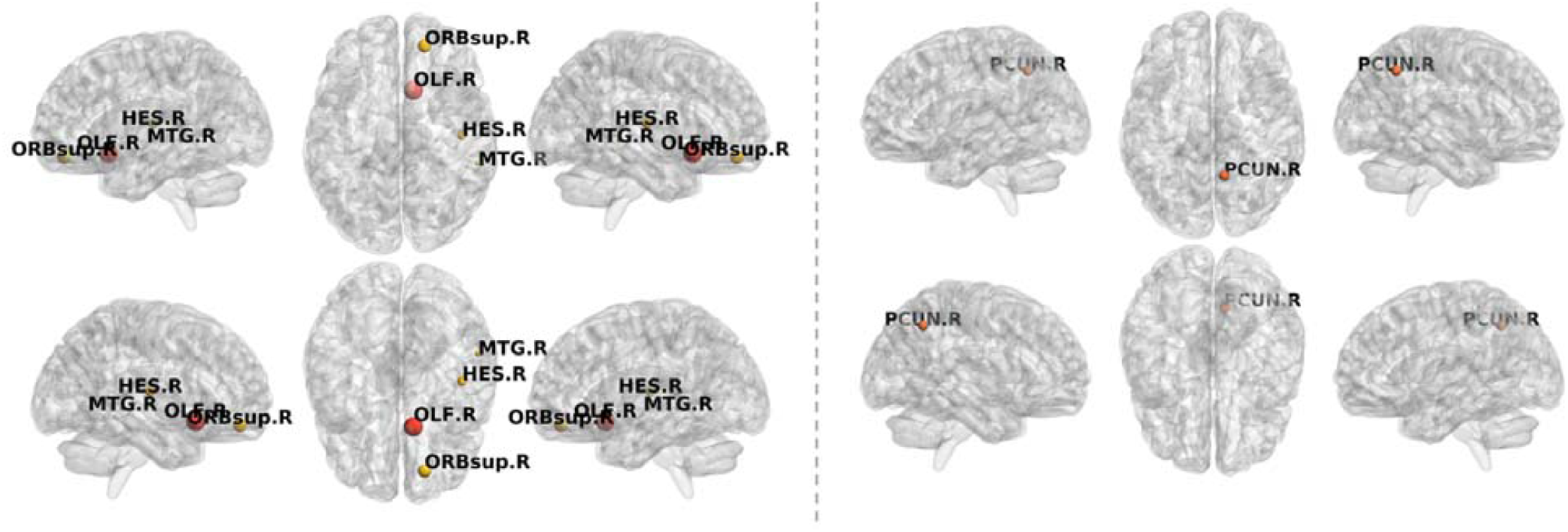
Node level graph metrics. Nodes with significant increases in NSPL in ADHD-I compared to ADHD-C (orange) and in ADHD-I compared to ADHD-C (red) are plotted in the left. Nodes with significant increases in betweenness centrality in ADHD-I compared to ADHD-C are plotted on the right.

Our results also show that the NSPL of the right heschl’s gyrus and right medial temporal gyrus is increased in the ADHD-I group compared to TDC. The NSPL of the olfactory cortex was increased in ADHD-I compared to ADHD-C (Fig.7A). Deficits in olfactory function are found in neurodegenerative and neuropsychiatric disorders and represent a topic of interest in ADHD [48]. Increased NSPL was found in the right olfactory region in ADHD-I compared to ADHD-C suggesting lesser integration. Deficits in olfactory ability have been linked to impulsive tendencies within the healthy population and have discriminatory features in identifying people at risk of impulse-control-related problems, supporting the planning of early clinical interventions [49]. Further studies are required to investigate whether functional network topology can be used as a biological marker for early diagnosis, treatment and prognosis of ADHD.

It is important to note that the proposed method measures non-linear relationships while correlation methods measure linear relationships, which may have resulted in the lower performance of correlation in inferring non-linear brain dynamics. Although PC performed relatively similarly to BrainNET in the simulation data, it didn’t achieve statistical significance in the ADHD data. This may be due to the false positives identified reducing the statistical power of the analysis. BrainNET can be added to the standard inference methods such as PC and correlation methods, by using a mask derived from BrainNET importance matrix and applying to the correlation matrix. The output from this combined method will have nodes determined by BrainNET, with Pearson correlation values assigned between the connections. This will avoid using arbitrary thresholds, increase the specificity of the standard inference methods by adding non-linearity and allowing analysis of connectivity changes between nodes, which cannot be performed with an adjacency matrix derived only from BrainNET.

#### Limitations

BrainNET takes relatively longer to infer the adjacency matrix than the correlation method. BrainNET took approximately 3 seconds per subject whereas the correlation and partial correlation method just took 0.001 and 9.3 seconds respectively. Longer running time makes BrainNET challenging to apply for voxel-wise analysis.

## 5. Conclusion

We describe BrainNET, a new network inference method to estimate fMRI connectivity that was adapted from Gene Regulatory methods. We validated the proposed model on ground truth simulation data [8]. BrainNET outperformed Pearson correlation in terms of accuracy and sensitivity across simulations and various confounders such as the presence of cyclic connections, and even with truncated fMRI sessions of only 2.5 min. We evaluated the performance of BrainNET on the open-source “ADHD 200 preprocessed” data from Neuro Bureau. BrainNET was able to identify significant changes in global graph metrics between ADHD groups and TDC, whereas correlation and PC was unable to find any differences. BrainNET can be used independently or combined with other existing methods as a useful tool to understand network changes and to determine the true network topology of the brain under various conditions and disease states.

## Acknowledgments

This research did not receive any specific grant from funding agencies in the public, commercial, or not-for-profit sectors.

## Author Disclosure Statement

No competing financial interests exist

## Author contributions

Gowtham Krishnan Murugesan conceptualized and designed the work, analyzed and interpreted the data, and wrote the paper. Dr. Joseph Maldjian and Dr. Kim Won Hwa provided expert knowledge and mentorship to develop the method. Ben Wagner contributed in developing fmri analysis. Chandan Ganesh,, Sahil Nalawade, and Dr. Elizabeth Davenport contributed to review the paper

---

**Supplementary Table 1:** Simulations and corresponding confounders.

| Simulation | Nodes | fMRI<br>session<br>length (min) | TR | Noise<br>(%) | HRF std<br>dev (s) | Other factors |
| --- | --- | --- | --- | --- | --- | --- |
| 1 | 5 | 10 | 3 | 1 | 0.5 |  |
| 2 | 10 | 10 | 3 | 1 | 0.5 |  |
| 3 | 15 | 10 | 3 | 1 | 0.5 |  |
| 4 | 50 | 10 | 3 | 1 | 0.5 |  |
| 5 | 5 | 60 | 3 | 1 | 0.5 |  |
| 6 | 10 | 60 | 3 | 1 | 0.5 |  |
| 7 | 5 | 250 | 3 | 1 | 0.5 |  |
| 8 | 5 | 10 | 3 | 1 | 0.5 | shared inputs |
| 9 | 5 | 250 | 3 | 1 | 0.5 | shared inputs |
| 10 | 5 | 10 | 3 | 1 | 0.5 | global mean confound |
| 11 | 10 | 10 | 3 | 1 | 0.5 | bad ROIs (time series<br>mixed with each other) |
| 12 | 10 | 10 | 3 | 1 | 0.5 | bad ROIs (new random<br>time series mixed in) |
| 13 | 5 | 10 | 3 | 1 | 0.5 | backwards connections |
| 14 | 5 | 10 | 3 | 1 | 0.5 | cyclic connections |
| 15 | 5 | 10 | 3 | 0.1 | 0.5 | stronger connections |
| 16 | 5 | 10 | 3 | 1 | 0.5 | more connections |
| 17 | 10 | 10 | 3 | 0.1 | 0.5 |  |
| 18 | 5 | 10 | 3 | 1 | 0 |  |
| 19 | 5 | 10 | 0.25 | 0.1 | 0.5 | neural lag = 100 ms |
| 20 | 5 | 10 | 0.25 | 0.1 | 0 | neural lag = 100 ms |
| 21 | 5 | 10 | 3 | 1 | 0.5 | 2-group test |
| 22 | 5 | 10 | 3 | 0.1 | 0.5 | Non stationary connection strength |
| 23 | 5 | 10 | 3 | 0.1 | 0.5 | stationary connection strength |
| 24 | 5 | 10 | 3 | 0.1 | 0.5 | only one strong<br>external input |
| 25 | 5 | 5 | 3 | 1 | 0.5 |  |
| 26 | 5 | 2.5 | 3 | 1 | 0.5 |  |
| 27 | 5 | 2.5 | 3 | 0.1 | 0.5 |  |
| 28 | 5 | 5 | 3 | 0.1 | 0.5 |  |

**Supplementary Table 2.** Sensitivity, Specificity, Accuracy, and C-sensitivity for each inference method across simulations. Optimum thresholds (with highest performance calculated using simulated data) were used for PC and correlation. The BrainNet threshold was set to [1/number of nodes]. No thresholds were needed for C-sensitivity. C-sensitivity is a measure of fraction of true positives that are estimated with a higher connection strength than the 95th percentile of the false positive distribution.

| Simulation | Sensitivity |  |  | Specificity |  |  | Accuracy |  |  | C-Sensitivity |  |  |
| --- | --- | --- | --- | --- | --- | --- | --- | --- | --- | --- | --- | --- |
|  | Corr | PC | BN | Corr | PC | BN | Corr | PC | BN | Corr | PC | BN |
| 1 | 0.97 | 1.00 | 0.99 | 0.96 | 0.90 | 0.89 | 0.97 | 0.98 | 0.97 | 0.85 | 0.98 | 0.90 |
| 2 | 0.94 | 0.99 | 0.96 | 0.99 | 0.87 | 0.93 | 0.94 | 0.98 | 0.96 | 0.48 | 0.96 | 0.90 |
| 3 | 0.95 | 1.00 | 0.95 | 0.95 | 0.80 | 0.92 | 0.95 | 0.98 | 0.95 | 0.68 | 0.92 | 0.87 |
| 4 | 1.00 | 1.00 | 0.92 | 0.68 | 0.59 | 0.95 | 0.99 | 0.99 | 0.92 | 0.91 | 0.88 | 0.90 |
| 5 | 0.96 | 1.00 | 1.00 | 1.00 | 1.00 | 0.97 | 0.97 | 1.00 | 0.99 | 1.00 | 1.00 | 1.00 |
| 6 | 0.99 | 1.00 | 1.00 | 1.00 | 1.00 | 0.99 | 0.99 | 1.00 | 1.00 | 1.00 | 1.00 | 1.00 |
| 7 | 1.00 | 1.00 | 1.00 | 1.00 | 1.00 | 0.97 | 1.00 | 1.00 | 0.99 | 1.00 | 1.00 | 1.00 |
| 8 | 0.75 | 0.90 | 0.95 | 1.00 | 0.90 | 0.81 | 0.80 | 0.90 | 0.93 | 0.06 | 0.47 | 0.67 |
| 9 | 0.75 | 0.90 | 0.96 | 1.00 | 1.00 | 0.88 | 0.80 | 0.92 | 0.95 | 0.00 | 0.20 | 0.78 |
| 10 | 0.78 | 0.99 | 0.98 | 1.00 | 1.00 | 0.94 | 0.83 | 0.99 | 0.97 | 0.91 | 1.00 | 0.94 |
| 11 | 0.80 | 0.93 | 0.93 | 1.00 | 0.79 | 0.73 | 0.82 | 0.91 | 0.91 | 0.18 | 0.01 | 0.12 |
| 12 | 0.93 | 0.99 | 0.96 | 0.95 | 0.77 | 0.93 | 0.93 | 0.96 | 0.95 | 0.74 | 0.84 | 0.86 |
| 13 | 1.00 | 1.00 | 0.95 | 0.00 | 0.20 | 0.86 | 0.80 | 0.84 | 0.93 | 0.24 | 0.24 | 0.72 |
| 14 | 0.79 | 0.93 | 0.95 | 1.00 | 1.00 | 0.88 | 0.82 | 0.94 | 0.94 | 0.10 | 0.35 | 0.48 |
| 15 | 0.75 | 1.00 | 0.98 | 1.00 | 1.00 | 0.86 | 0.80 | 1.00 | 0.95 | 0.65 | 1.00 | 0.87 |
| 16 | 0.94 | 1.00 | 0.98 | 0.96 | 0.86 | 0.75 | 0.95 | 0.96 | 0.92 | 0.42 | 0.81 | 0.81 |
| 17 | 0.92 | 0.99 | 0.98 | 1.00 | 0.99 | 0.95 | 0.93 | 0.99 | 0.98 | 0.69 | 0.99 | 0.96 |
| 18 | 0.99 | 1.00 | 0.99 | 1.00 | 0.94 | 0.91 | 0.99 | 0.99 | 0.98 | 1.00 | 1.00 | 0.91 |
| 19 | 0.85 | 1.00 | 1.00 | 0.80 | 0.80 | 0.91 | 0.84 | 0.96 | 0.98 | 0.80 | 0.80 | 0.96 |
| 20 | 0.85 | 1.00 | 1.00 | 0.80 | 0.80 | 0.91 | 0.84 | 0.96 | 0.98 | 0.80 | 0.80 | 0.98 |
| 21 | 0.97 | 1.00 | 0.98 | 0.96 | 0.90 | 0.89 | 0.97 | 0.98 | 0.96 | 0.85 | 0.98 | 0.88 |
| 22 | 0.99 | 0.95 | 0.99 | 0.20 | 0.31 | 0.84 | 0.83 | 0.82 | 0.96 | 0.20 | 0.26 | 0.81 |
| 23 | 0.75 | 0.96 | 0.92 | 1.00 | 1.00 | 0.88 | 0.80 | 0.97 | 0.91 | 0.60 | 1.00 | 0.63 |
| 24 | 0.75 | 0.90 | 0.85 | 1.00 | 1.00 | 0.79 | 0.80 | 0.92 | 0.84 | 0.57 | 0.82 | 0.36 |
| 25 | 0.92 | 0.98 | 0.94 | 0.84 | 0.75 | 0.83 | 0.91 | 0.93 | 0.92 | 0.44 | 0.71 | 0.72 |
| 26 | 0.98 | 1.00 | 0.93 | 0.35 | 0.41 | 0.79 | 0.86 | 0.88 | 0.90 | 0.55 | 0.59 | 0.61 |
| 27 | 0.90 | 0.96 | 0.96 | 0.85 | 0.80 | 0.82 | 0.89 | 0.93 | 0.93 | 0.59 | 0.76 | 0.76 |
| 28 | 0.92 | 0.98 | 0.98 | 0.90 | 0.89 | 0.87 | 0.91 | 0.96 | 0.95 | 0.43 | 0.84 | 0.88 |
| Average | 0.90 | 0.98 | 0.96 | 0.86 | 0.83 | 0.88 | 0.89 | 0.95 | 0.95 | 0.60 | 0.76 | 0.80 |

## Notes

#### Summary of Updates

Added results from Partial correlation and extended analysis of ADHD

## References

[1] O. Sporns, “Graph theory methods: applications in brain networks,” Dialogues in Clinical Neuroscience, vol. 20, no. 2, p. 111, 2018.

[2] A. Fornito, A. Zalesky, and M. Breakspear, “The connectomics of brain disorders,” Nat Rev Neurosci, vol. 16, no. 3, pp. 159–72, Mar 2015, doi: 10.1038/nrn3901.

[3] A. Avena-Koenigsberger, B. Misic, and O. Sporns, “Communication dynamics in complex brain networks,” Nature Reviews Neuroscience, vol. 19, no. 1, p. 17, 2018.

[4] R. L. Buckner et al., “Cortical hubs revealed by intrinsic functional connectivity: mapping, assessment of stability, and relation to Alzheimer’s disease,” Journal of neuroscience, vol. 29, no. 6, pp. 1860–1873, 2009.

[5] N. A. Crossley et al., “The hubs of the human connectome are generally implicated in the anatomy of brain disorders,” Brain, vol. 137, no. 8, pp. 2382–2395, 2014.

[6] D. E. Warren et al., “Network measures predict neuropsychological outcome after brain injury,” Proceedings of the National Academy of Sciences, vol. 111, no. 39, pp. 14247–14252, 2014.

[7] A. Fornito, A. Zalesky, and E. Bullmore, Fundamentals of brain network analysis. Academic Press, 2016.

[8] S. M. Smith et al., “Network modelling methods for FMRI,” Neuroimage, vol. 54, no. 2, pp. 875–91, Jan 15 2011, doi: 10.1016/j.neuroimage.2010.08.063.

[9] G. Zaharchuk, E. Gong, M. Wintermark, D. Rubin, and C. Langlotz, “Deep learning in neuroradiology,” American Journal of Neuroradiology, vol. 39, no. 10, pp. 1776–1784, 2018.

[10] G. Murugesan et al., “Single Season Changes in Resting State Network Power and the Connectivity between Regions: Distinguish Head Impact Exposure Level in High School and Youth Football Players,” Proc SPIE Int Soc Opt Eng, vol. 10575, Feb 2018, doi: 10.1117/12.2293199.

[11] G. Murugesan et al., “Changes in resting state MRI networks from a single season of football distinguishes controls, low, and high head impact exposure,” Proc IEEE Int Symp Biomed Imaging, vol. 2017, pp. 464–467, Apr 2017, doi: 10.1109/ISBI.2017.7950561.

[12] B. Saghafi et al., “Quantifying the Association between White Matter Integrity Changes and Subconcussive Head Impact Exposure from a Single Season of Youth and High School Football using 3D Convolutional Neural Networks,” Proc SPIE Int Soc Opt Eng, vol. 10575, Feb 2018, doi: 10.1117/12.2293023.

[13] N. Williams and R. N. Henson, “Recent advances in functional neuroimaging analysis for cognitive neuroscience,” ed: SAGE Publications Sage UK: London, England, 2018.

[14] E. Pellegrini et al., “Machine learning of neuroimaging to diagnose cognitive impairment and dementia: a systematic review and comparative analysis,” arXiv preprint 1804.01961, 2018.

[15] T. J. O’Neill, E. M. Davenport, G. Murugesan, A. Montillo, and J. A. Maldjian, “Applications of resting state functional mr imaging to traumatic brain injury,” Neuroimaging Clinics, vol. 27, no. 4, pp. 685–696, 2017.

[16] T. Turki, J. T. Wang, and I. Rajikhan, “Inferring gene regulatory networks by combining supervised and unsupervised methods,” in 2016 15th IEEE International Conference on Machine Learning and Applications (ICMLA), 2016: IEEE, pp. 140–145.

[17] D. M. Camacho, K. M. Collins, R. K. Powers, J. C. Costello, and J. J. Collins, “Next-generation machine learning for biological networks,” Cell, 2018.

[18] J. D. Finkle, J. J. Wu, and N. Bagheri, “Windowed Granger causal inference strategy improves discovery of gene regulatory networks,” Proceedings of the National Academy of Sciences, vol. 115, no. 9, pp. 2252–2257, 2018.

[19] A. Irrthum, L. Wehenkel, and P. Geurts, “Inferring regulatory networks from expression data using tree-based methods,” PloS one, vol. 5, no. 9, p. e12776, 2010.

[20] K. Hilger and C. J. Fiebach, “ADHD symptoms are associated with the modular structure of intrinsic brain networks in a representative sample of healthy adults,” Network Neuroscience, vol. 3, no. 2, pp. 567–588, 2019.

[21] P. Lin et al., “Global and local brain network reorganization in attention-deficit/hyperactivity disorder,” Brain imaging and behavior, vol. 8, no. 4, pp. 558–569, 2014.

[22] F. Saeed, “Towards quantifying psychiatric diagnosis using machine learning algorithms and big fMRI data,” Big Data Analytics, vol. 3, no. 1, p. 7, 2018.

[23] S. Cortese et al., “Toward systems neuroscience of ADHD: a meta-analysis of 55 fMRI studies,” American Journal of Psychiatry, vol. 169, no. 10, pp. 1038–1055, 2012.

[24] J. Sidlauskaite, K. Caeyenberghs, E. Sonuga-Barke, H. Roeyers, and J. R. Wiersema, “Whole-brain structural topology in adult attention-deficit/hyperactivity disorder: Preserved global–disturbed local network organization,” NeuroImage: Clinical, vol. 9, pp. 506–512, 2015.

[25] P. Bellec, C. Chu, F. Chouinard-Decorte, Y. Benhajali, D. S. Margulies, and R. C. Craddock, “The neuro bureau ADHD-200 preprocessed repository,” Neuroimage, vol. 144, pp. 275–286, 2017.

[26] M. P. Milham, D. Fair, M. Mennes, and S. H. Mostofsky, “The ADHD-200 consortium: a model to advance the translational potential of neuroimaging in clinical neuroscience,” Frontiers in systems neuroscience, vol. 6, p. 62, 2012.

[27] N. Tzourio-Mazoyer et al., “Automated anatomical labeling of activations in SPM using a macroscopic anatomical parcellation of the MNI MRI single-subject brain,” Neuroimage, vol. 15, no. 1, pp. 273–289, 2002.

[28] A. Abraham et al., “Machine learning for neuroimaging with scikit-learn,” Frontiers in neuroinformatics, vol. 8, p. 14, 2014.

[29] C. J. Stam and J. C. Reijneveld, “Graph theoretical analysis of complex networks in the brain,” Nonlinear biomedical physics, vol. 1, no. 1, p. 3, 2007.

[30] L. Breiman, Classification and regression trees. Routledge, 2017.

[31] F. Petralia, P. Wang, J. Yang, and Z. Tu, “Integrative random forest for gene regulatory network inference,” Bioinformatics, vol. 31, no. 12, pp. i197–i205, 2015.

[32] C. Strobl, A.-L. Boulesteix, A. Zeileis, and T. Hothorn, “Bias in random forest variable importance measures: Illustrations, sources and a solution,” BMC bioinformatics, vol. 8, no. 1, p. 25, 2007.

[33] A. Yamashita et al., “Harmonization of resting-state functional MRI data across multiple imaging sites via the separation of site differences into sampling bias and measurement bias,” vol. 17, no. 4, p. e3000042, 2019.

[34] J.-P. Fortin et al., “Harmonization of cortical thickness measurements across scanners and sites,” vol. 167, pp. 104–120, 2018.

[35] J. Wang, X. Wang, M. Xia, X. Liao, A. Evans, and Y. J. F. i. h. n. He, “GRETNA: a graph theoretical network analysis toolbox for imaging connectomics,” vol. 9, p. 386, 2015.

[36] A. Hagberg et al., “Networkx. High productivity software for complex networks,” Webová strá nka https://networkx.lanl.gov/wiki, 2013.

[37] J. Wang, X. Zuo, and Y. J. F. i. s. n. He, “Graph-based network analysis of resting-state functional MRI,” vol. 4, p. 16, 2010.

[38] S. Achard and E. J. P. c. b. Bullmore, “Efficiency and cost of economical brain functional networks,” vol. 3, no. 2, p. e17, 2007.

[39] P. Geurts, D. Ernst, and L. Wehenkel, “Extremely randomized trees,” Machine learning, vol. 63, no. 1, pp. 3–42, 2006.

[40] M. Wehenkel, C. Bastin, C. Phillips, and P. Geurts, “Tree ensemble methods and parcelling to identify brain areas related to Alzheimer’s disease,” in 2017 International Workshop on Pattern Recognition in Neuroimaging (PRNI), 2017: IEEE, pp. 1–4.

[41] K. Supekar, M. Musen, and V. Menon, “Development of large-scale functional brain networks in children,” PLoS biology, vol. 7, no. 7, p. e1000157, 2009.

[42] S. Achard and E. Bullmore, “Efficiency and cost of economical brain functional networks,” PLoS computational biology, vol. 3, no. 2, p. e17, 2007.

[43] X. Qian et al., “Large-scale brain functional network topology disruptions underlie symptom heterogeneity in children with attention-deficit/hyperactivity disorder,” vol. 21, p. 101600, 2019.

[44] A. D. Barber et al., “Connectivity supporting attention in children with attention deficit hyperactivity disorder,” vol. 7, pp. 68–81, 2015.

[45] A. F. J. T. J. o. p. Arnsten, “The emerging neurobiology of attention deficit hyperactivity disorder: the key role of the prefrontal association cortex,” vol. 154, no. 5, p. I, 2009.

[46] A. dos Santos Siqueira, B. Junior, C. Eduardo, W. E. Comfort, L. A. Rohde, and J. R. Sato, “Abnormal functional resting-state networks in ADHD: graph theory and pattern recognition analysis of fMRI data,” BioMed Research International, vol. 2014, 2014.

[47] L. Weyandt, A. Swentosky, and B. G. J. D. n. Gudmundsdottir, “Neuroimaging and ADHD: fMRI, PET, DTI findings, and methodological limitations,” vol. 38, no. 4, pp. 211–225, 2013.

[48] A. Ghanizadeh, M. Bahrani, R. Miri, and A. Sahraian, “Smell identification function in children with attention deficit hyperactivity disorder,” Psychiatry investigation, vol. 9, no. 2, p. 150, 2012.

[49] A. M. Herman, H. Critchley, and T. J. S. r. Duka, “Decreased olfactory discrimination is associated with impulsivity in healthy volunteers,” vol. 8, no. 1, p. 15584, 2018.

